# Circumstances and factors of sleep-related sudden infancy deaths in Japan

**DOI:** 10.1101/2020.05.04.076612

**Authors:** Motoki Osawa, Yasuhiro Ueno, Noriaki Ikeda, Kazuya Ikematsu, Takuma Yamamoto, Wataru Irie, Shuji Kozawa, Hirokazu Kotani, Hideki Hamayasu, Takehiko Murase, Keita Shingu, Marie Sugimoto, Ryoko Nagao, Yu Kakimoto

## Abstract

**Background:** Sudden unexpected death in infancy (SUDI) comprises both natural and unnatural causes of death. However, a few epidemiological surveys have investigated SUDI in Japan.

**Objective:** This retrospective study was conducted to investigate the recent trends of circumstances and risk factors of sleep-related SUDI cases.

**Methods:** Forensic pathology sections from eight universities participated in the selection of subjects from 2013 to 2018. Data obtained from the checklist form were analyzed based on information at postmortem.

**Results:** There were 259 SUDI cases consisting of 145 male infants and 114 female infants with a mean birth weight of 2888 ± 553 and 2750 ± 370 g, respectively. Deaths most frequently occurred among infants at 1 month of age (18%). According to population-based analyses, the odds ratio (95% confidence interval) of mother’s age ≤19 years was 11.1 (6.9–17.7) compared with ages 30–39. The odds ratio for the fourth- and later born infants was 5.2 (3.4–7.9) compared with the frequency of first-born infants. The most frequent time of day for discovery was between 7 and 8 o’clock. Co-sleeping was recorded for 61%, and the prone position was found for 40% of cases at discovery. Mother’s smoking habit exhibited an adds ratio of 4.5 (2.9–5.8).

**Conclusion:** This study confirmed the trends that have been observed for sudden infant death syndrome; particularly, very high odds ratios were evident for teenage mothers and later birth order in comparison with those in other developed countries. The child of a young mother tended to die within 2 months of age. To our knowledge, this is the first report of an extensive survey of sleep-related SUDI in Japan.

## Introduction

Sudden infant death syndrome (SIDS) is the possible cause-of-death for sudden infant death during sleep, in which all known identifiable conditions that might engender sudden and unexpected death must be excluded by postmortem examinations. However, several pathologists have changed their diagnostic preferences since 2004 primarily because of the difficult distinction of SIDS from accidental asphyxia or natural diseases such as arrhythmias and metabolic disorders. [1-3] Reluctance to use the term has decreased the number globally over the years. [4-6] In Japan, around 500 cases of SIDS were recorded in the 1990s, but recent diagnostic numbers have decreased to fewer than 100 cases per year. [7]

Currently, another broad term has become popular, i.e., sudden unexpected death in infancy (SUDI) or sudden unexpected infant death (SUID). Although SUDI/SUID originally has been used as an umbrella term for the initial presentation of explained or unexplained infant deaths, it is interpreted to include several categories such as SIDS (R95), ill-defined and unknown cause of mortality (R99), and accidental suffocation or strangulation in bed (W75). [8] Several attempts have been made to categorize SUDI/SUID, but the terminology and classification have not yet been defined clearly. [9]

In recent years, a protocol of investigation items has been officially standardized. [10,11] For instance, in the U.S., the Centers for Disease Control and Prevention published guidelines and a reporting form, which is designated as the SUID Investigation Reporting Form. [12] In Japan, a list of items to be investigated is used as a checklist form in cases of infant death. [13] A system is in operation for clinicians and pathologists to ascertain circumstances and to investigate background factors. Information from death scene investigation (DSI) acquired by experts is indispensable to fill out the form. [14] Furthermore, the maternity passbook, which records information about the mother and the child during pregnancy as well as after childbirth, is beneficial. We previously analyzed forensic autopsy cases of sudden infant deaths after vaccination using this passbook. [15]

While the childcare environment differs according to region and time, there are few epidemiological data describing infant deaths during sleep in Japan. [16,17] This population-based retrospective study was conducted to investigate the recent trends of sleep-related SUDI using the checklist form at multiple centers.

## Methods

Deaths of sleep-related SUDI cases were extracted from autopsy files for the period of 6 years from 2013 to 2018. Inclusion criteria for cases were age not more than 12 months, and the collapse occurring during sleep in an unexpected manner. Based on the cause and manner of death, infant deaths were grouped as follows: (1) infants who died of SIDS, (2) infants who died of other natural diseases, (3) infants who died of accidental injuries, (4) infants who died of non-accidental injuries, and (5) infants with unexplained manner of death. [18] The term ‘unexplained’ implies insufficient evidence of the causative disease or event. [9] In this study, we selected cases of groups (1) and (5) and those of suspected accidental suffocation during sleep.

Postmortem examinations included histology, toxicology, biochemistry, virology, and bacteriology. [19,20] Tests for assessing inherited metabolic disorders were conducted nationwide in the routine examination for newborns. [21] Genetic testing for arrhythmic disorders was also conducted for cases examined in this study. [22] Data used for analysis consisted of DSI information, therapeutic information in emergency care, and maternity passbook. The checklist form, consisting of 41 items, was filled in initially by each center. Then the lists were transferred to one site to confirm unclear issues and aggregate the data.

The forensic pathology sections of the following eight universities participated in this study: Kitasato University School of Medicine, Mie University School of Medicine, Kyoto University Graduate School of Medicine, Hyogo College of Medicine, Kobe University Graduate School of Medicine, Graduate School of Medical Science Kyushu University, Graduate School of Biomedical Sciences Nagasaki University, and Tokai University School of Medicine. The areas of these facilities cover six prefectures in which approximately 14% of the entire population resides, and this percentage was applied to calculate the annual rate per 1000. Every sudden infant death had been autopsied, but there could have been some deaths that had not received autopsy outside major centers in Japan, whose exact number remained unknown. Although DSI was performed by the police for all cases, not all items were optimal, particularly, for the sleep environment such as sleep surface, wrapping, and clothing. The principal investigator obtained approval for this retrospective study from the Institutional Review Board for Clinical Research, Tokai University. This study was also approved by the respective ethical committees of the faculties as a collaborative study.

The original causes of death (*n* = 259) were SIDS and suspicious SIDS in 94 cases (36%), unexplained in 75 cases (29%), potential asphyxia in bed in 51 cases (20%), and other causes in 39 cases (15%). We investigated each candidate case carefully at a meeting, and selected subjects in which terminal events remained at speculation irrespective of the diagnosis in the death certificate. Suspected causes of asphyxia were supposed to be due to accidental overlay and swallowing in bed. Inflammation of the airway, including bronchitis, accounted for 22 cases, comprising the largest group among “others.” Despite of such histological evidence, the pathologists reconsidered that these cases might also be regarded as sleep-related SUDI because of co-sleeping, in which coexistent factors may have served as contributors in causing death.

The statistical data of live births recorded during 2013–2018 in the Japanese population at the National Institute of Population and Social Security Research (http://www.ipss.go.jp/p-info/) were used as the control. Prevalence information of regular tobacco consumption was available at Japan Tobacco Inc. as a questionnaire survey performed in 2016 (https://www.jti.co.jp/investors/press_releases/2016/0728_01.html). The numbers of smokers and nonsmokers in the 20s and 30s of female volunteers were used for the control.

Logistic regression analyses based on population data were performed to determine the associations of independent variables as estimated using the odds ratio and confidence interval (CI). For significant differences, the *p* value was calculated using chi-square tests. Statistical analyses were conducted using BellCurve in Excel, ver. 3.20 (Social Survey Research Information Co., Tokyo, Japan).

## Results

A total of 259 cases were collected at multiple centers for the 6-year period. The circumstances and factors were investigated using the checklist form filled with information at postmortem.

The annual frequency of sleep-related SUDI was estimated to approximately 0.3 per 1000 births. However, the value could be slightly underestimated because of the possibility that some cases could have been out of management by these facilities.

Table 1 presents the number of subjects according to sex, birth weight, gestation week, maternal age, parity and maternal smoking habit along with the population data. Table 2 summarizes the odds ratios (95% CI) and *p* values in terms of the related factors. The factors were also analyzed for two groups, i.e., the early group in which death occurred within 2 months of age (*n* = 98) and the late group in which death occurred after 3 months (*n* = 161).

**Table 1.**
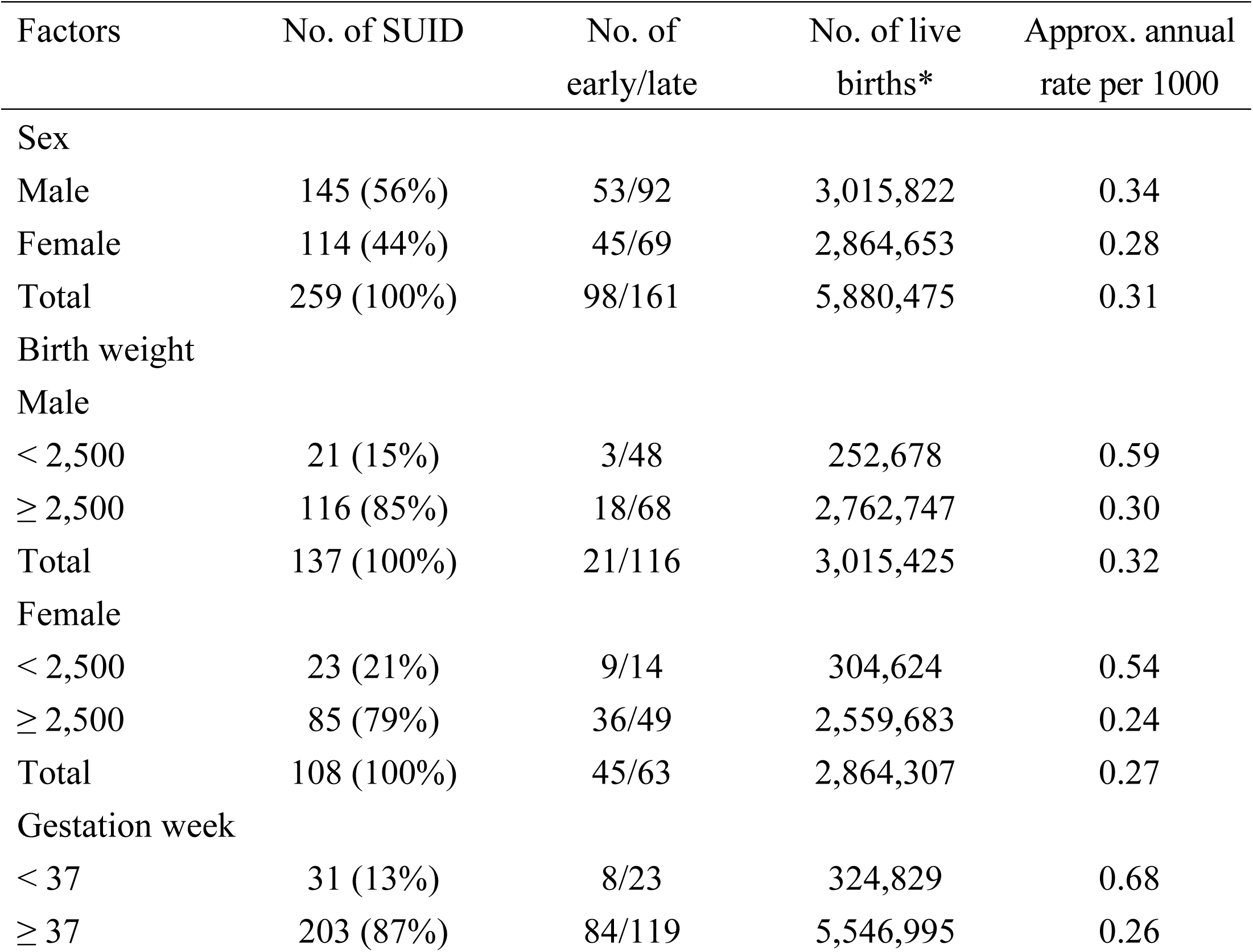
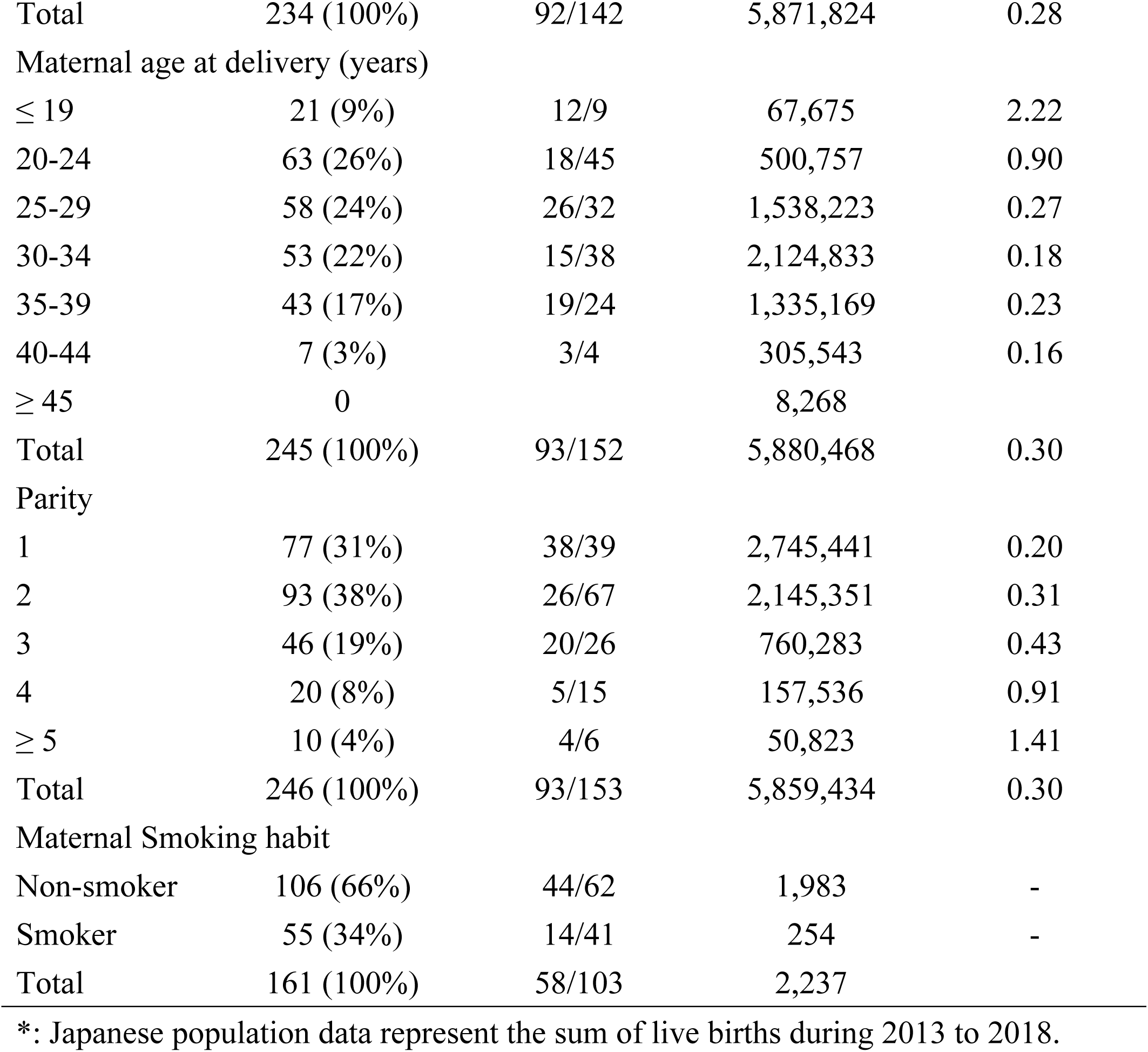
Number of infants of SUID subjects and live births

**Table 2.** Odds ratios (95% CI) and *p* values of SUID including early and late occurrences

| Factors | Total (0–11 month) | Early (0–2 month) | Late (3–11 month) |
| --- | --- | --- | --- |
| Sex |  |  |  |
| Male | 1.2 (0.9–1.5), *p* = 0.13 | 1.1 (0.8–1.7), *p* = 0.58 | 1.3 (0.9–1.7), *p* = 0.14 |
| Female (ref. group) | 1.0 | 1.0 | 1.0 |
| Birth weight |  |  |  |
| Male |  |  |  |
| < 2,500 | 2.0 (1.3–3.2), *p* = 0.003 | 0.7 (0.2–2.2), *p* = 0.52 | 2.9 (1.7–4.9), *p* < 0.001 |
| ≥ 2,500 (ref. group) | 1.0 | 1.0 | 1.0 |
| Female |  |  |  |
| < 2,500 | 2.3 (1.4–3.6), *p* < 0.001 | 2.1 (1.0–4.4), *p* = 0.042 | 2.4 (1.3–4.3), *p* = 0.003 |
| ≥ 2,500 (ref. group) | 1.0 | 1.0 | 1.0 |
| Gestation week |  |  |  |
| < 37 | 2.6 (1.8–3.8), $p < 0.001$ | 1.6 (0.8–3.4), $p = 0.18$ | 3.3 (2.1–5.2), $p < 0.001$ |
| ≥ 37 (ref. group) | 1.0 | 1.0 | 1.0 |
| Maternal age at delivery (years) |  |  |  |
| ≤ 19 | 11.1 (6.9–17.7), $p < 0.001$ | 18.0 (9.3–34.8), $p < 0.001$ | 7.4 (3.7–14.9), $p < 0.001$ |
| 20–29 | 2.1 (1.6–2.8), $p < 0.001$ | 2.2 (1.4–3.5), $p < 0.001$ | 2.1 (1.5–2.9), $p < 0.001$ |
| 30–39 (ref. group) | 1.0 | 1.0 | 1.0 |
| ≥ 40 | 0.8 (0.4–1.7), $p = 0.56$ | 1.0 (0.3–3.2), $p = 0.96$ | 0.7 (0.3–2.0), $p = 0.51$ |
| Parity |  |  |  |
| 1 (ref. group) | 1.0 | 1.0 | 1.0 |
| 2–3 | 1.7 (1.3–2.3), $p < 0.001$ | 1.1 (0.7–1.7), $p = 0.61$ | 2.3 (1.6–3.3), $p < 0.001$ |
| ≥ 4 | 5.1 (3.4–7.8), $p < 0.001$ | 3.1 (1.5–6.5), $p = 0.0012$ | 7.1 (4.2–12.1), $p < 0.001$ |
| Maternal smoking habit |  |  |  |
| Non-smoker (ref. group) | 1.0 | 1.0 | 1.0 |
| Smoker | 4.1 (2.9–5.8), $p < 0.001$ | 2.5 (1.3–4.6), $p = 0.002$ | 5.2 (3.4–7.8), $p < 0.001$ |

### Age

Fig 1 shows the age distribution at the time of death. It was observed that deaths most frequently occurred in infants at 1 month of age, consisting of 45 cases (18%). The number was found to decrease with age. Deaths occurring within 6 months after birth accounted for 180 cases (72%).

**Fig 1.**
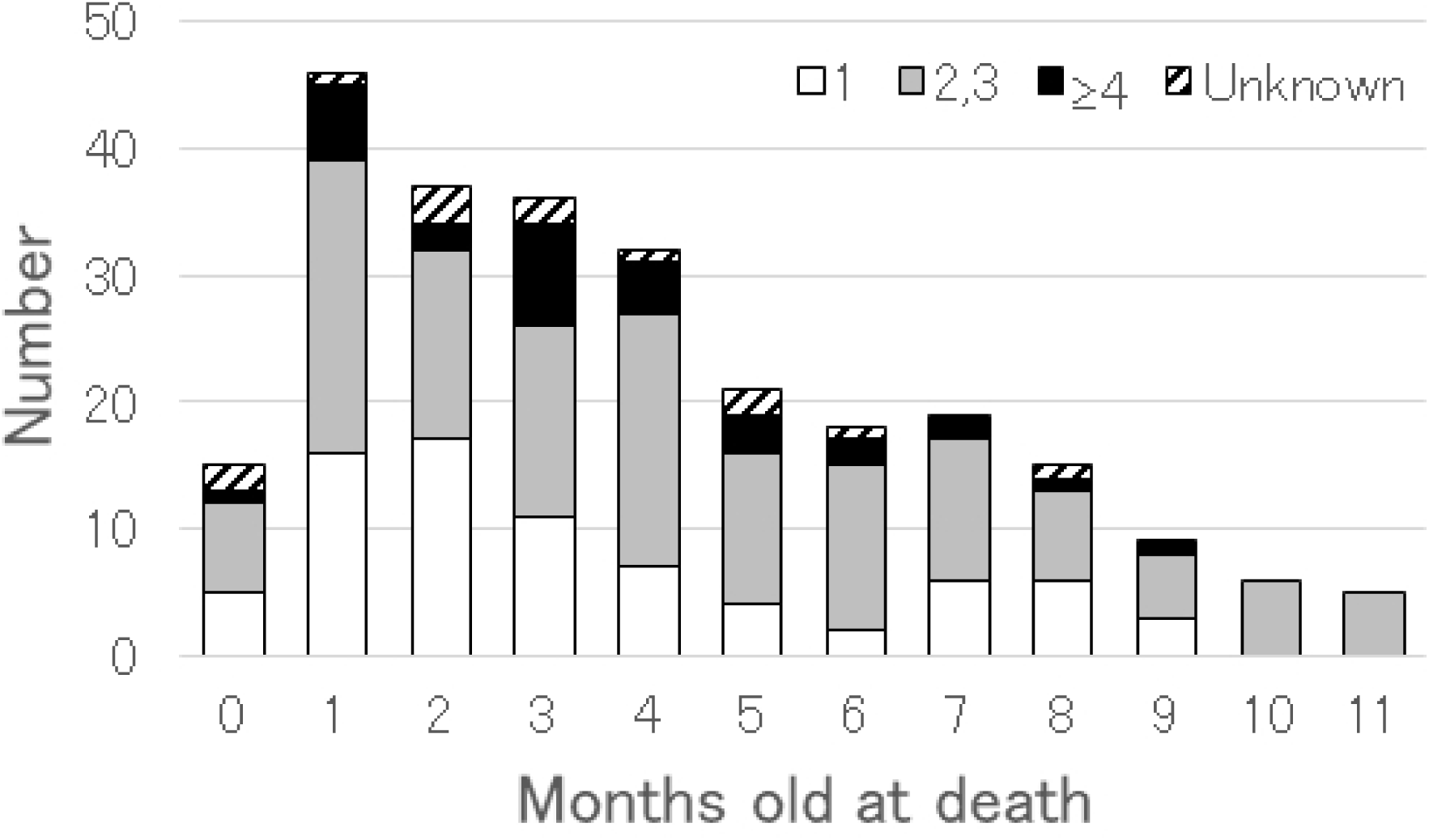
Age distribution of sleep-related SUDI infants (*n* = 259) examined in this study. Key indicates the birth order of infants, in which the column is divided into four groups of the first-born infants (blank), the second- and third-born ones (gray), more than the fourth-born ones (black), and unknown ones (diagonal).

### Birth weight and gestation weeks

The mean (± S.D.) birth weight of SUDI infants was 2885 ± 556 g for male subjects and 2763 ± 466 g for female subjects. These birth weights were lower by 191 g (6%) and 227 g (8%), respectively, than the national mean birth weights of 3076 g for males and 2990 g for females recorded in 2017. The percentage of low birth weight infants was significantly higher than the control group in both sexes. For low birth weight infants, the highest odds ratio of 2.9 was observed in the late male group.

Infants of premature birth were found to be 2.6 times more likely to die from SUDI than those of mature birth. The late group also showed higher odds ratio than the early group.

### Maternal age and birth order

The odds ratio of incidence of infant death of mothers whose age ≤19 years was the highest at 11.1 compared with mothers aged 30–39 years, and that for mothers aged 20–29 years was 2.1, which showed significant differences. Furthermore, the child of a teenage mother tended to die within 2 months of age, compared with other generations (12/21 to 81/225, *p* = 0.04). This finding indicated that mothers of a younger age, especially teenage, should be considered as the most important risk factor for the occurrence of sleep-related SUDI.

In terms of the birth-order distribution, there were 31% of first-born infants, 38% of second-born infants, 19% of third-born infants, 8% of fourth-born infants, 2% (6 cases) of fifth-born infants, 1% (3 cases) of sixth-born infants, and 0.4% (1 case) of seventh-born infant. The odds ratio to the fatal frequency among the first-born infants clearly indicated that later birth order constituted an important risk factor. Moreover, as shown in Figure 1, there were more first-born infants in the early group (38/92) than in the late group (39/154) (*p* < 0.001).

### Time of discovery and sleeping position

Fig 2A shows the distribution of the time of day when an unresponsive infant was found. There were 30 cases (12%) found between 7 and 8 o’clock a.m., which was the most frequent time. A large peak was evident between 6 and 9 o’clock in the morning.

**Fig 2.**
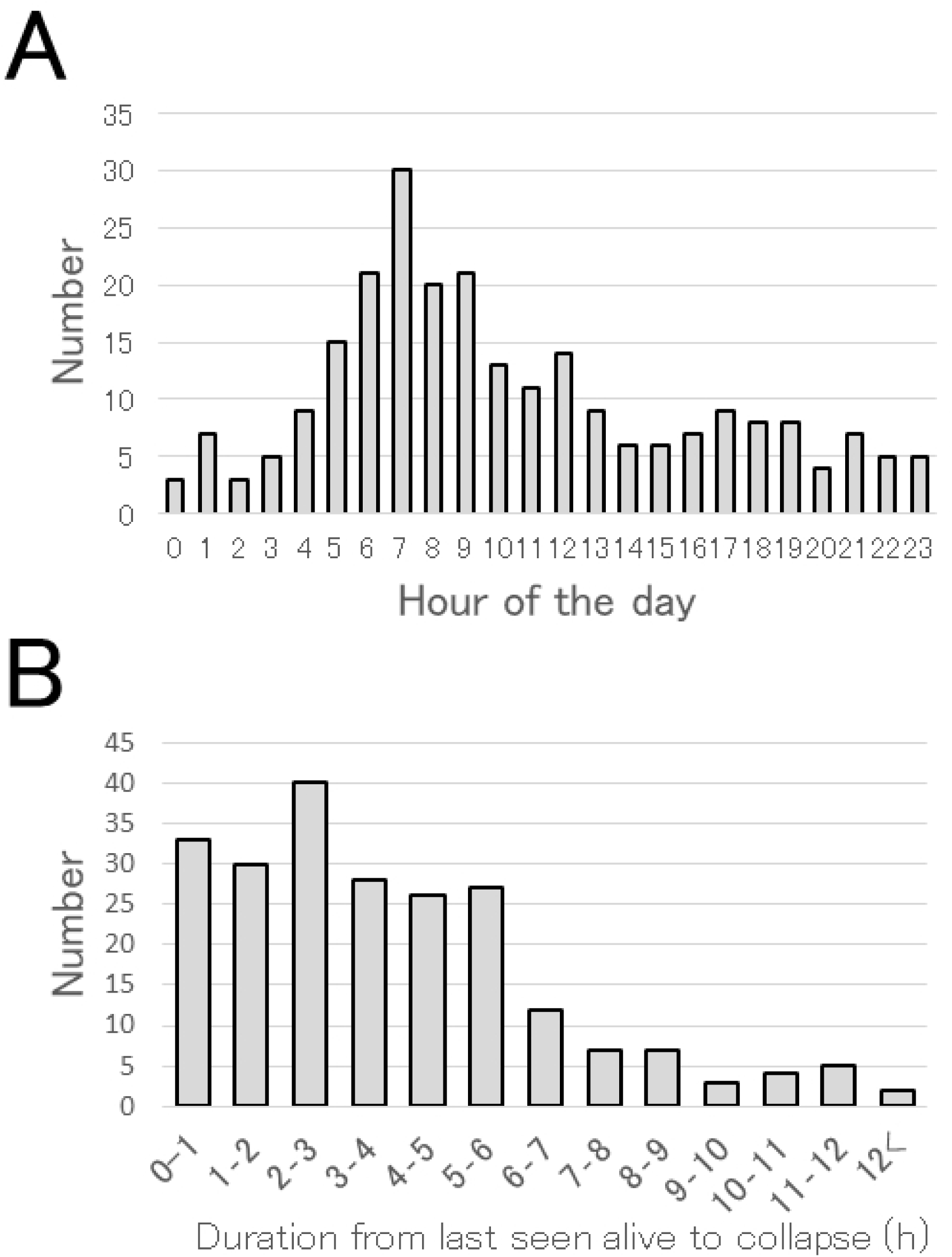
Time of discovery. A. Distribution of the time of the day at which the first responder found an unresponsive infant (*n* = 246); B. the time difference between the time the infant was last seen alive and first found deceased (*n* = 222).

Fig 2B depicts that the duration between the last time the infant was found alive and the time of discovery of being unresponsive (*n* = 222), which varied widely from approximately 10 min to 13 h. The mean duration was 4.1±2.7 h. The collapse was discovered within 6 h in the majority of cases (*n* = 184, 83%).

The first responder (*n* = 252), who discovered the unresponsive infant, was the mother in 188 cases (75%), followed by the father in 49 cases (19%), a grandmother in 8 cases (3%), a childminder in 2 cases (1%), and others in 6 cases (2%).

Co-sleeping was recorded for 143 cases (61%) among a total of 230 available cases. Table 3 presents the child sleeping position when the collapse was discovered. The prone position in late SUDI infants accounted for 52% of cases, and there were 19% of cases with the prone position in the early group.

**Table 3.** Sleep position at the scene

|  | Total (0–11 month) | Early (0–2 month) | Late (3–11 month) |
| --- | --- | --- | --- |
| Spine | 117 (52%) | 60 (72%) | 57 (40%) |
| Prone | 89 (40%) | 16 (19%) | 73 (52%) |
| Side | 16 (7%) | 7 (9%) | 9 (6%) |
| Others | 2 (1%) | 0 | 2 (1%) |
| Total | 224 (100%) | 83 (100%) | 141 (100%) |

### Maternal smoking habit

The descriptions in the maternity passbook entries are considered to reflect the smoking habits before and during the early phase of pregnancy. We attempted to obtain the smoking rate of the mothers of SUDI cases. A significant risk of SUDI was evident with an odds ratio of 4.5 compared with the general rate. Although there were limited cases wherein the information related to the number of cigarettes (*n* = 30) was available, the mean number was 11 cigarettes/day.

## Discussion

Results of the present investigation of sleep-related SUDI cases were consistent with risk factors such the smoking habit of parents in the large epidemiological surveys of SIDS. [23] However, some differences were evident. The peak age of death is generally 2 months in SIDS surveys, [4,24] but that among the present SUDI infants was 1 month of age. A particularly higher risk was observed among teenage mothers than that found in an earlier study. [25] Moreover, the mothers tended to lose their infants at age 0–2 months.

The most frequent birth order associated with infant death due to sleep-related SUDI was the second birth order. Blair et al. [26] reported that SIDS was most frequent among first-born children in the UK, although it was earlier presumed to be frequent in large families. Data from Taiwan indicate that the first-, second-, and third- and later born children account for 36%, 40%, and 24% of SIDS, respectively. [27] The distribution in the present study was more similar to that reported in Taiwan.

Traditional bedding of cotton mat, known as futon, on the floor is common in Japan. Therefore, it is more appropriate to use the term co-sleeping (sharing a sleeping surface) than bed-sharing. Co-sleeping is a common style of sleep, and Tokutake et al. [17] reported that 84% of mothers practice co-sleeping, of whom half also practice breastfeeding. The father was found to be the first responder in up to 20% of cases, and in most of these cases the father also co-slept and discovered the infant death upon awakening. The risk of SIDS among infants who co-sleep was found to be significantly high in several earlier studies. [28,29] Nevertheless, the effects of co-sleeping on the occurrence of SUDI, if any, could not be evaluated in this study because of the absence of good control subjects.

It is a traditional practice in Japan to lay infants in the supine position. However, 42% of infants were found in the prone position, of which frequency was higher than that reported in an earlier study. [17] Li et al. [30] reported that 60% of SIDS infants were found in the prone position in the United States. It is possible that turning over by infants during sleep is a causal factor. However, approximately 30% of infants in the early group who were unable to turn over were found in the prone and side positions. They might have been placed in the prone position or been breastfed during co-sleeping, but the original position was not recorded sufficiently at DSI.

A relationship between the occurrence of SIDS and the smoking habit of parents has been found in Japan. [31] In the present investigation, the incidence of pregnant mother’s smoking among SUDI cases was 34%. This incidence in the general female population was reported as 11%, which also resulted in the high odds ratio of 4.5 in this study. According to Anderson et al., [32] the incidence of SUDI more than doubles when a parent is smoking during the period of pregnancy. The odds ratio increases along with the number of cigarettes up to 20. It is evident that infants co-sleeping with someone who smokes exhibit the highest risk for SUID. [33]

Pasquale-Styles et al. [34] reported that asphyxia and suffocation occur more than presumed in many situations such as bed-sharing, overlay, wedging, prone position, obstruction of the nose and mouth, and coverage of the head. Postmortem findings alone are not generally sufficient to explain the cause of these deaths. Consequently, the diagnoses often lack consistency. [3,4,7] In addition, there exists a difficulty of the current situation in Japan, particularly in DSI that is performed by police officers who are not well trained. Garstang et al. [24] indicated that police-led DSI does not comply with practical information. After a new law related to child health was enacted in 2018, child death reviews will be introduced to the society in the near future. These reviews in combination with multiple agencies will be helpful in investigating the sleeping environments of infants in detail.

In conclusion, we conducted an effective epidemiological analysis of sleep-related SUDI using the official checklist form. This approach has revealed the present critical features prevailing in the country. This report is the first of an extensive study of SUDI in Japan.

## Acknowledgements

We thank Tomohiro Nozima for his extensive support like inputting the large data and mailing.

